# Gene transfer among viruses substantially contributes to gene gain of giant viruses

**DOI:** 10.1101/2023.09.26.559659

**Authors:** Junyi Wu, Lingjie Meng, Morgan Gaïa, Hiroyuki Hikida, Yusuke Okazaki, Hisashi Endo, Hiroyuki Ogata

## Abstract

Horizontal gene transfers (HGTs) integrate all forms of life and viruses into a vast network of gene flow, which facilitates the transmission of genes beyond vertical inheritance and enhances genomic evolution. HGT is known to occur between closely related viruses. We hypothesized that there is frequent HGT among nucleocytoviruses, a group of diverse but evolutionarily related DNA viruses encoding hundreds to thousands of genes. However, the frequency of viral HGT (vHGT) has not been systematically investigated for nucleocytoviruses. We reconciled over 4,700 gene trees with a robust viral species tree that contains 195 reference viral genomes mainly from cultivation as a reference to infer evolutionary events, including gene gains (gene duplication, origination, and vHGT) and losses. The inferred evolutionary scenarios revealed that the genomes of these viruses have undergone numerous gene gain and loss events, with vHGT representing 28% to 42% of gene gain events in each viral order. By integrating the evolutionary paths of multiple viruses, our data suggest that vHGT is crucial for nucleocytovirus evolution.

## Introduction

The viral phylum *Nucleocytoviricota*^1^ encompasses diverse large and giant double-stranded DNA viruses that infect a wide spectrum of eukaryotes, from protists to mammals. Some of the giant viruses possess hundreds to thousands of genes^2,3^. By encoding numerous unknown or uncharacterized genes, nucleocytoviruses challenge the conventional “Gene Pickpocket Hypothesis^4^”, which postulates that viruses largely acquire new genes from their host cellular organisms through horizontal gene transfer (HGT). Although many phylogenetic gene trees support HGT from cellular organisms^5-7^, up to 70% of the nucleocytovirus genes are unique to the phylum and lack detectable homologs in the cellular world. In the Gene Pickpocket Hypothesis, the lack of cellular homologs of these viral genes has been accounted by high mutation rates or relaxed functional constraints that erased trackable ancestral signals^8,9^. However, the genes present only in nucleocytoviruses show regular features in their sequence evolution without markedly high substitution rates^10^, calling into question the possibility that they were acquired from cellular hosts.

In addition to HGT from cellular organisms, gene duplication has also been suggested as a crucial mechanism driving gene gain in nucleocytovirus evolution. Suhre found that one-third of the genes in mimivirus have at least one paralog in its genome^11^. During a continuous passage experiment of vaccinia virus in HeLa cells, genes subject to selection pressure were found to duplicate, thereby providing immediate fitness advantages within a few passages^12^. These examples underscore the importance of gene duplication in the expansion of the gene repertoire of nucleocytoviruses during their evolution. Gene gain can also possibly occur through *de novo* creation. Previous studies have shown such mechanisms for cellular organisms, with the progressive constitution of proto-genes to protein-encoding genes through successive mutations and selection^13,14^. Assuming the infected cell is part of the virus life cycle, the virocell^15^, then the same mechanisms may also occur for viral genomes during their replication. This could partially explain the high proportion of unique genes in nucleocytoviruses, as proposed for pandoraviruses^16^.

Another possible evolutionary mechanism for gene gain in nucleocytoviruses is virus-to-virus HGT (vHGT). vHGT is considered as one of the main driving forces in the evolution of phages^17-19^ and some animal infected viruses^20^ (e.g., Influenza viruses). Only a few cases have been reported on the existence of vHGT in nucleocytoviruses so far. For instance, the phyletic patterns of gene presence and absence suggested potential vHGT events between the mimivirus and marseillevirus lineages through the co-infection of the two viruses in the same amoeba cell^21^. Other evidence emerged from the observation of inconsistencies between gene trees and the species tree; virmyosin^22^ and viractin^23^ genes were transferred among viruses of the *Imitervirales* order and a gene of unknown function was transferred between pandoraviruses and mollivirus^24^. These cases suggest the importance of vHGT in these viruses. However, previous systematic evolutionary reconstructions of nucleocytovirus genomes used methods that do not distinguish different mechanisms of gene gains^25,26^, which further humper the investigation of the impact of vHGT on nucleocytovirus evolution compared with other gene gain mechanisms.

To assess the frequency of vHGT during nucleocytoviruses evolution, we used the amalgamated likelihood estimation^27^ (ALE) method, which is a tree reconciliation method that compares a gene tree to a species tree to identify different evolutionary events. ALE is a probabilistic approach that supports the detection of three classes of gene gain (gene duplication, HGT, and origination) and gene loss. We applied the ALE method on the gene families and viral species tree for nucleocytoviruses. In this framework, the HGT events inferred by ALE are likely to be vHGT, while the origination events represent either HGT from outside the tree, such as cellular organisms or other viruses not included in our dataset, or *de novo* gene creation. In this study, we generated a robust viral tree (“species tree”) for 195 reference nucleocytoviruses (mostly cultured ones) and performed the reconciliation of over 4,700 gene families to infer the evolutionary events. The analyses revealed massive gene gain and loss events across different viral orders. vHGT was found to represent 28% to 42% of gene gain events that were inferred to have occurred during the evolution of viral orders. These data highlight the previously overlooked contribution of vHGT in nucleocytoviruses evolution.

## Results

### Robust viral tree for *Nucleocytoviricota*

A prerequisite for tree reconciliation by ALE for evolutionary inference is a robust phylogenetic tree for viruses that reflects their speciation events. By using seven concatenated marker genes that are considered important for viral replication and display consistent phylogenetic signals^1^, we reconstructed a phylogenetic tree of nucleocytoviruses (hereafter called the viral tree). After applying an iterative process to remove, by manual inspection, viruses that contribute to unstable branches, such as medusavirus and Heterosigma akashiwo virus, we obtained the final viral tree (Fig. 1). In this viral tree, all branches were supported (UFboot > 95% or SH-aLRT > 80%) except one deep branch (the position of the recently proposed viral order *Pandoravirales*^1^). The viral tree contained 195 nucleocytoviruses that span 6 viral orders: the *Chitovirales* and the *Asfuvirales* orders of the *Pokkesviricetes* class, and the *Imitervirales, Algavirales, Pandoravirales*, and *Pimascovirales* orders of the *Megaviricetes* class. The viral tree was rooted between the two class, following the official International Committee on Taxonomy of Viruses taxonomy^1^ (Fig. 1). This viral tree was then used as the reference for tree reconciliation of individual gene trees for different gene families (orthogroups; OGs), which allows for the inference of different evolutionary events (gene gains and losses).

**Figure 1.**
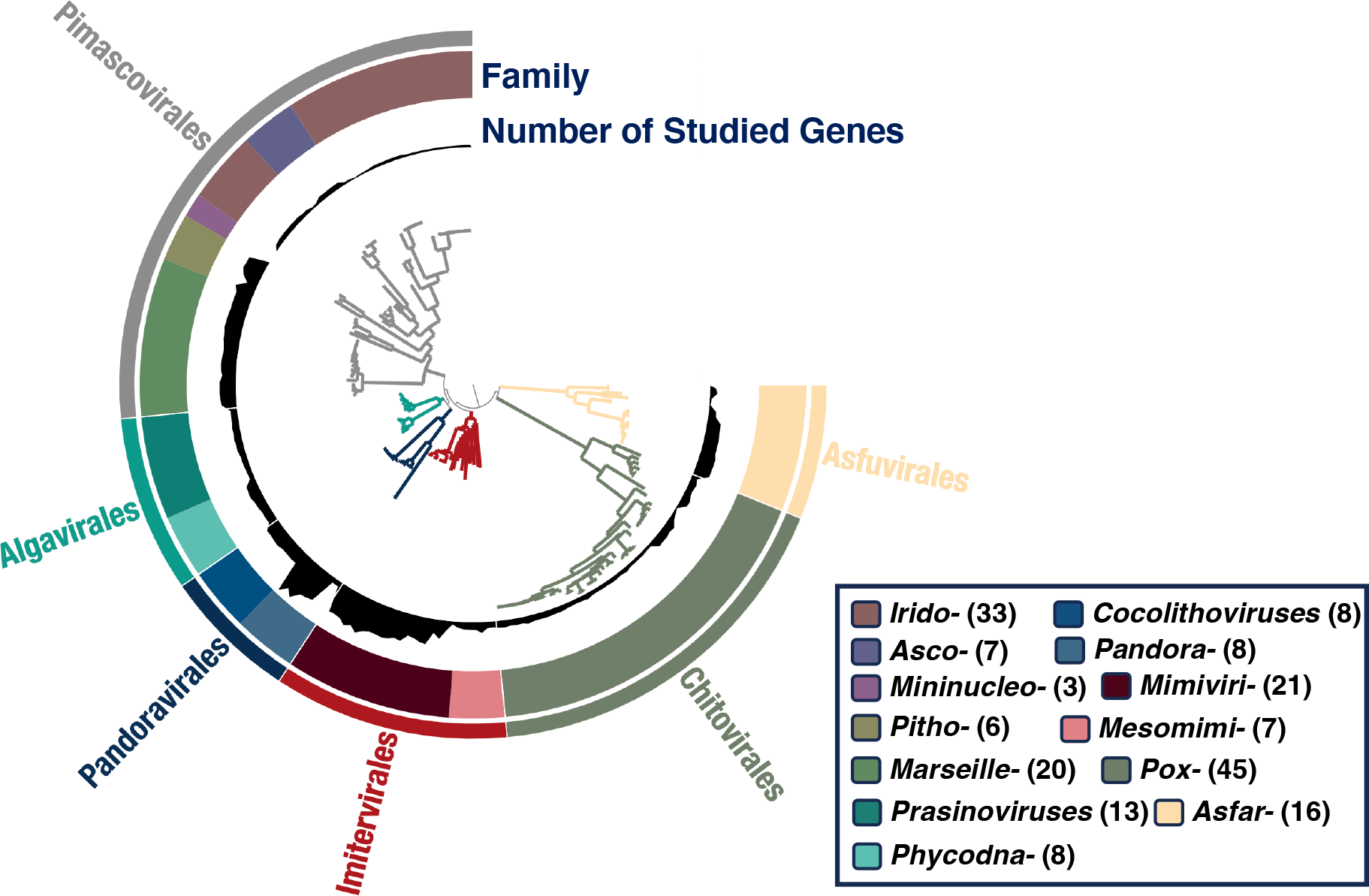
Viral tree of *Nucleocytoviricota*. The phylogenetic tree is based on concatenated seven marker genes and constructed using IQ-TREE with the Q.pfam+F+l+l+R8+C60 model. The outer layer designates the viral orders, the middle layer represents viral families, and the inner layer indicates the number of genes (ranging from 11 to 1,207) that were successfully inferred for tree reconciliation analysis. The number in parentheses in the legend panel indicates the number of viruses in each family and “-” is the abbreviation of “viridae”.

### Gene trees for reconciliation

Predicted genes in the 195 viral genomes were grouped into 8,876 OGs, excluding singletons. Of these, a total of 4,786 OGs met the requirements for confident ALE reconciliations, and evolutionary scenarios involving gene gain (gene duplication, vHGT, and origination) and/or gene loss were inferred for them (Supplementary Table 1 for example, see Methods). These OGs cover an average of 74.3% of genes from individual viral genomes (Fig. S1).

### The compensation of massive gene loss by gene gain

Reconciliation of 4,786 gene trees with the reference viral tree allowed us to infer 19,442 gene gain and 16,999 gene loss events, along with a conservative estimate of the number of genes at ancestral nodes (Fig. 2). The number of genes in the ancestral genomes of viral orders ranged from 53 for *Chitovirales* to 191 for *Imitervirales*. The ancestor of the class *Pokkeviricetes* was inferred to encode at least 42 genes, while the ancestor of *Megaviricetes* had 113 genes. The ancestral genome at the root of *Nucleocytoviricota* was inferred to encode at least 40 genes, which is consistent with previous estimations^25,28^. The number of gene loss events (average 45.3%) was comparable to the number of gene gain events (average 54.7%) over the course of evolution irrespective of viral orders (Fig. 2). For example, the extant mimivirus encodes 805 genes considered for the evolutionary reconciliation. Over the course of the evolution from the root of *Nucleocytoviricota* to the extant mimivirus, it was inferred that the virus acquired 1,553 genes and lost 683 genes (see Table S1 for other examples).

**Figure 2.**
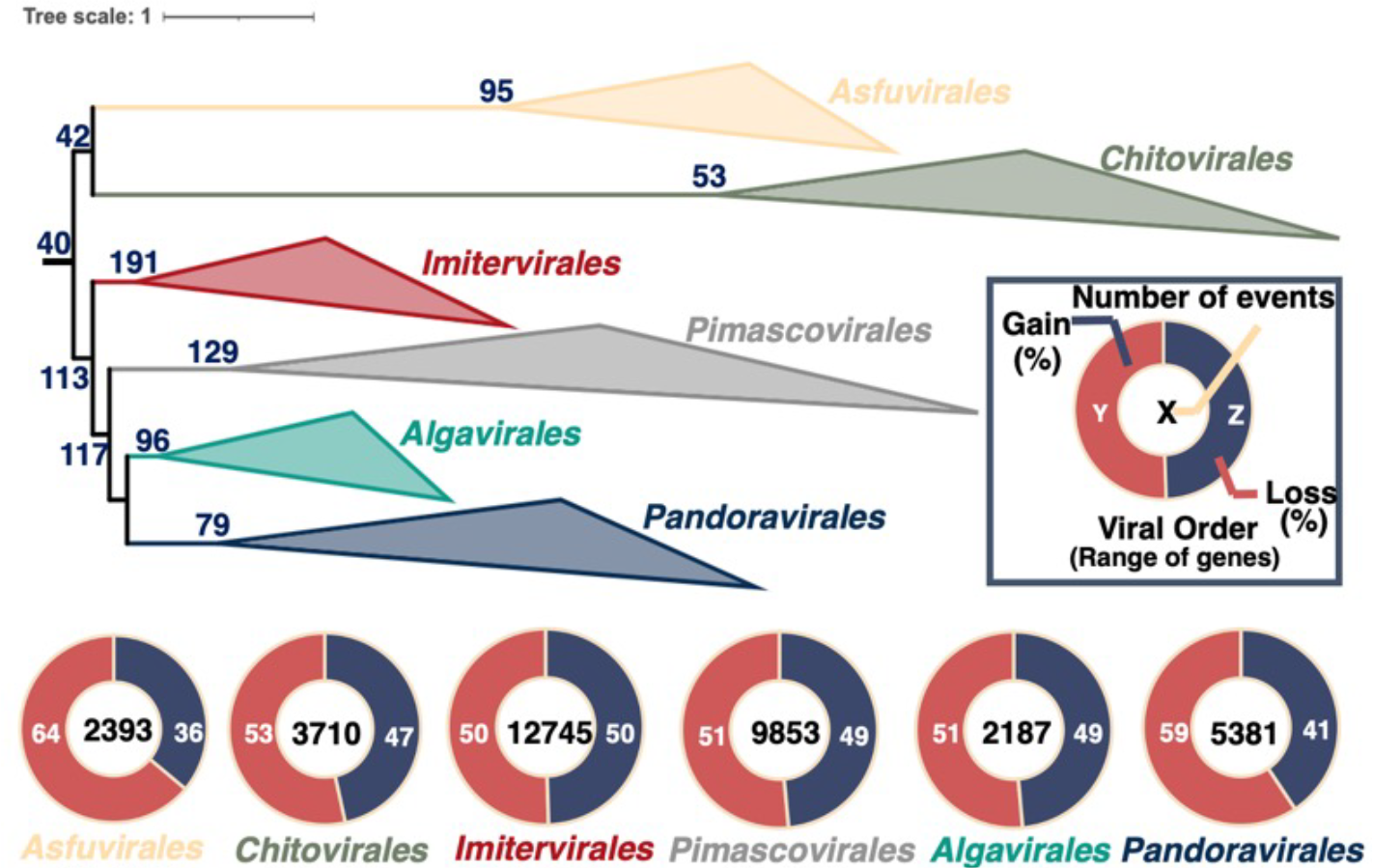
Evolutionary inference of gene gain and loss events for *Nucleocytoviricota*. The phylogenetic tree, collapsed at the level of viral orders, is displayed with the number of genes inferred to be present at the ancestral nodes. Accompanying this tree is a series of pie charts illustrating the proportion of gene gain and loss events, with the total number of events (gain + loss) given at the center of each chart.

### vHGT contributes to gene gains at a comparable level as other two mechanisms

The ALE method can help distinguish three mechanisms of gene gain: gene duplication, origination (HGT from outside the tree of *Nucleocytoviricota*, such as cellular organisms or other viruses, or *de novo* gene creation), and vHGT. Our evolutionary reconstruction revealed that the contributions of these three mechanisms to the genome evolution were comparable with each other (Fig. 3). The contribution of gene duplication to gene gain varied from 26% (*Asfuvirales*) to 47% (*lmitervirales*), origination accounted for from 18% (*Imitervirales*) to 37% (*Pandoravirales*), and the contribution of vHGT was from 28% (*Pandoravirales*) to 42% (*Asfuvirales* and *Pimascovirales*).

**Figure 3.**
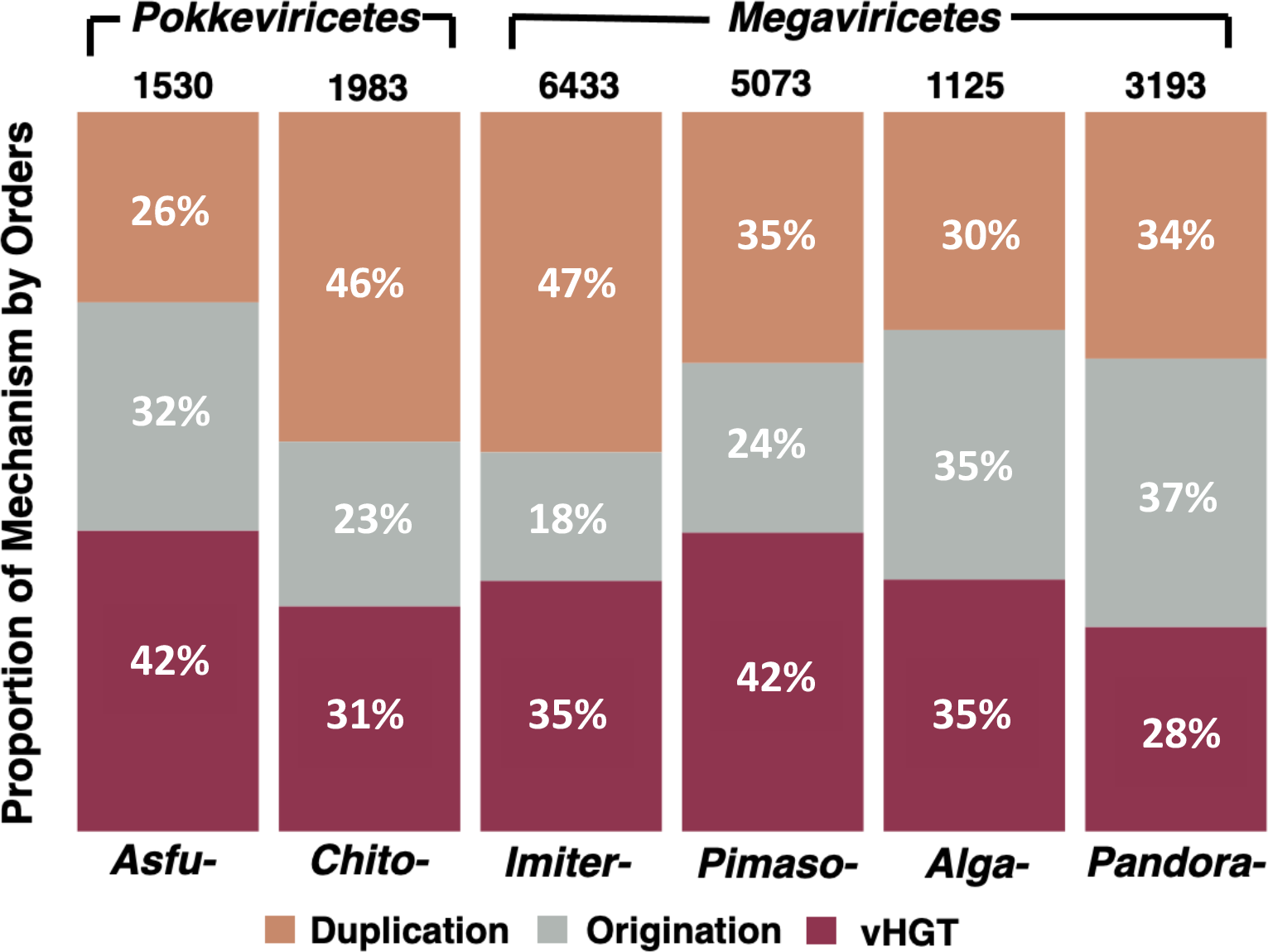
Contributions of different gene gain mechanisms. This bar plot depicts the contribution of each gene gain mechanism (duplication, origination, and vHGT) with the total number of gene gains indicated above each bar.

The above approach, however, could possibly artificially recognize parallel acquisitions of a cellular gene (or a gene homologous to a cellular gene) by multiple nucleocytoviruses as vHGT. To exclude this possibility for the contribution of vHGT, we focused on a subset of the 4,786 OGs that were absent in cellular organisms and thus specific to viruses. Because parallel acquisitions by viruses from cellular organisms are unlikely for those OGs, we labelled these OGs as viral-specific OGs. There were 3,535 viral-specific OGs. The gain events of this subset again showed a similar level of vHGT contribution (27% to 43%) (Fig. S2). One example of the visualization of reconciliation from an OG is shown in Figure S3.

### Conservative estimates of the rates of gene repertoire changes

To investigate the timing of evolutionary events, we calculated evolutionary rates as the number of evolutionary events normalized to the length of the branch in which the events occurred. We first calculated relative evolutionary divergence (RED)^29^ values for the nodes of the viral tree. The RED value serves as a measure of divergence time, with ‘0’ corresponding to the root of the tree and ‘1’ corresponding to the leaf (extant viruses). When the rates of gene gain and loss were plotted against the RED value, there was a clear acceleration in the rates in the recent past (high rates near RED = 1) (Fig. 4a and Fig. S4a). Evolutionary rates (except originations) were significantly higher for the recent past (RED 2: 0.95) than for the period before (RED < 0.95) for gene gain, loss, duplication, and vHGT (Mann-Whitney U test, *P* < 0.001; Fig. 4b and Fig. S4b). The evolutionary inference of moumouvirus genomes, for example, depicts the high rates of recent evolution. Three moumouviruses had diverged in recent past (RED = 0.98), and their mean ANl (Average Nucleotide ldentity) was 84.7%. ln the period from RED = 0.98 to 1, they gained and lost a lot of genes: the average gene gain was 63 and the average gene loss was 162. These results suggest conservative estimates of evolutionary events for deeper branches compared with more recent ones, as expected due to the absence of extinct viruses in the analyzed dataset.

**Figure 4.**
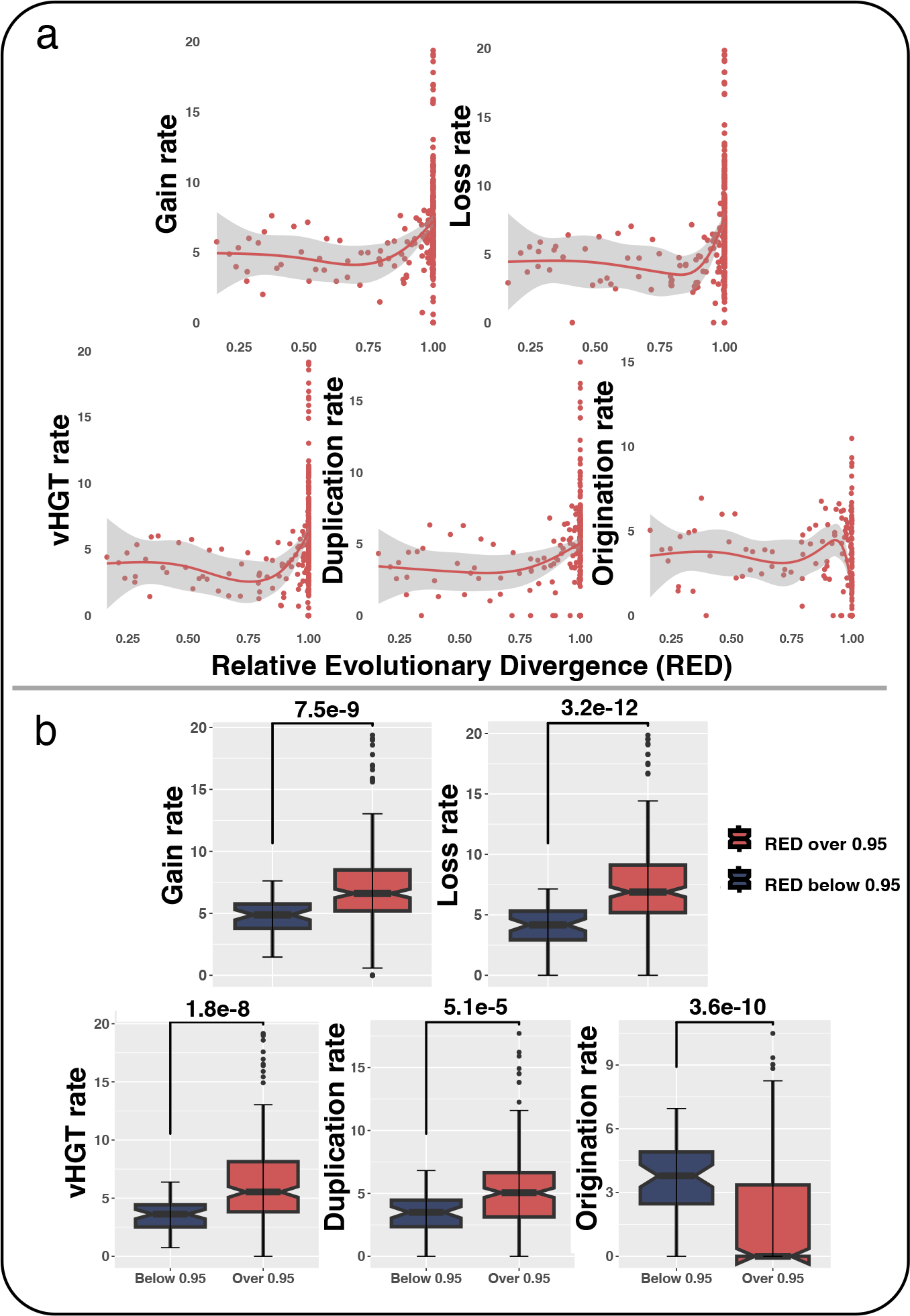
Rates of different evolutionary events along the divergence of *Megaviricetes*. (a) The evolutionary rates for different evolutionary events are plotted against the divergence measured by RED. (b) Boxplot provides a comparison of evolutionary rates between recent (RED ≥ 0.95) and earlier periods (RED < 0.95). *P*-values from Mann-Whitney U test are shown above the graph.

## Discussion

In the present study, we demonstrated that vHGT contributes to nucleocytovirus gene gains at a comparable level as gene duplication and origination (Fig. 3), regardless of whether all or viral-specific genes were analyzed (Fig S2). These results suggest that nucleocytoviruses can acquire genes from three principal genetic pools: genomes of their own, host organisms, and other related viruses. Their own genomic material supports the generation of new genes by gene duplication or *de novo* creation. The genetic pool of cellular organisms can also be accessed through HGT, potentially broadening their functional repertoires. Lastly, vHGT allows nucleocytoviruses to access the gene pool of other nucleocytoviruses. By bridging the gene pools of viruses, vHGT particularly allows interplay between the evolutionary paths of different viruses. The study of virmyosin^22^ detected in metagenomic assembled nucleocytoviruses presents a clear case of vHGT. The virmyosin gene was suggested to have originated in an HGT event from an ancient cellular organism to an ancient *Imitervirales* virus. Rather than maintaining this gene only through vertical transmission in a single viral lineage, the virmyosin gene was spread into different *Imitervirales* lineages via vHGT, leading to its presence across a diverse range of viruses. It is believed that after the ancestral virus acquired this gene’s beneficial functions, virmyosin was then transferred to other viruses, possibly during the co-infection of different viruses in the same host. Genes that are beneficial for one virus are likely to be useful for other viruses that infect the same host in the same environment. Our results on the high frequency of vHGT support this concept.

In the phylum *Nucleocytoviricota*, viruses showed broad but heterogenous host tropism^2^, suggesting frequent host switching during their evolution. We speculate that vHGT plays an important role in both adaptation to host (and surrounding environmental conditions) and host transition. The vHGT is likely to happen during the co-infection of the same host cell. Beneficial genes newly introduced by vHGT may contribute to different aspects in the viral infection cycle depending on their functions (i.e., from invasion into the host to release of virions). The transferred beneficial genes have experienced selection during donors’ evolution and thus potentially benefitable for the recipient virus. Thus, vHGT may enhance the adaptation of the recipient virus in the same host. In another scenario, where two viruses have overlapped host ranges, vHGT may unlock the ability of infecting a new host for both viruses, in a way analogous to one of the mechanisms of the zoonotic spillover^30^ (i.e., the transmission of viruses from wild animals to humans through an intermediate host). Therefore, the high frequency of vHGT may enhance nucleocytoviruses’ adaptation to a host and transition to a new host.

The ALE tree reconciliation method can systematically and quantitively identify evolutionary events, however some limitations still exist. Previous works^25,28^ and the current study identified only a small portion of genes as inherited from ancestral viruses (Fig. 2). However, the inferred number of genes in the ancestral nucleocytoviruses is likely a conservative estimate because of three limitations when inferring evolutionary events using the ALE tree reconciliation method. First, the inference is affected by the sampling of extant viruses. The virus genomes that we used in this study likely do not represent the actual virus diversity in nature. For example, the highly diverse nucleocytovirus genomes discovered from metagenomes^31,32^ were not considered in this study because of the incompleteness of their gene repertoires. The method likely missed the evolutionary events involving unsampled viruses (including many extinct viruses in deep branches). Second, genes that have been lost across all studied viruses cannot be incorporated into the evolutionary inference process. Third, for OGs that are specific to a viral order, we used the subtree of the viral tree corresponding to the order for reconciliation (see Methods). This may result in an underestimation of origination and loss events at deep branches older than the order formation (see Methods for details). Consequently, the number of genes inferred to have been present in ancestral nucleocytoviruses (Fig. 2) does not necessarily represent their gene content, but rather represents the lower limit of the number of genes encoded by these ancestral viruses. For example, the last common ancestor of *Imitervirales* was inferred to have possessed 191 genes, which represent genes successfully inherited by some of the extant *Imitervirales* members analyzed in this study. Therefore, our analysis provided conservative estimates for gene gain and loss events, especially in deeper branches.

The conservative estimation in evolutionary events partially explains the observed “J-shape” in our evolutionary rate reconstruction (i.e., low rates in deep branches; Fig.4a and S4a). The “J-shape” pattern can be interpreted in two ways. 1) recent acceleration of evolutionary rates or 2) lack of data for extinct viral species. The recent acceleration will lead to the evolutionary scenario where most lineages of the nucleocytoviruses become “giant” in recent period (i.e., RED close to 1). This interpretation is however unlikely due to the difficulty in explaining the sudden, recent and concomitant evolutionary paradigm shift for all nucleocytovirus lineages. It is more plausible that the real evolutionary rates in deeper branches were higher than estimated in this study and comparable to those in the recent past. Subsequently, the actual genomic content evolution of nucleocytoviruses through extensive gene gain and loss could have occurred since early stages of their evolution.

In this study, we used a probabilistic tree reconciliation tool to quantitatively infer four evolutionary events (i.e., gene duplication, vHGT, originations, and losses) along the evolution of nucleocytoviruses. We verified vHGT is as frequent as other gene gain events. We suggest that through vHGT, nucleocytoviruses can get access to other nucleocytoviruses’ gene repertoires and have chance to get beneficial genes that has experienced selection in the evolution of other viruses. We decided not to include Metagenome-Assembled Genomes (MAGs) due to possible contamination and incompleteness. The restricted diversity of genomes in our dataset hence resulted in robust yet conservative estimate of evolutionary events. Should the MAGs of nucleocytoviruses reach higher quality and completion, their inclusion would help reconstruct more concrete evolutionary scenarios. The frequent vHGT observed from this study coupled with its potential implications in facilitating transitions and adaptations to new hosts, lead us to posit an open question for future research: “To what extent does vHGT influence transitions to and adaptations in new hosts during the evolutionary path of nucleocytoviruses?” The challenge remains due to the high proportion of unknown function gene families and the lack of markers for host transition. We believe that with the improvement in deciphering gene functions in nucleocytoviruses (e.g., genes responsible for host tropism), this question can be solved.

## Methods

### Collection of reference and complete *Nucleocytoviricota* genomes

The reference genomes of viruses in the phylum *Nucleocytoviricota* were collected from the National Center for Biotechnology Information (NCBI) RefSeq databases by searching for the taxonomy *Nucleocytoviricota* or collected from published nucleocytoviruses isolation paper (Genomic data is available in www.genome.jp/ftp/db/community/vHGT/vHGT_data/). Our curated dataset includes 195 viruses that cover six proposed viral orders: *Algavirales, Asfuvirales, Chitovirales, Imitervirales, Pandoravirales*, and *Pimascovirales*.

### Reconstruction of the robust viral tree

Each virus in the dataset was subject to *de novo* protein sequence prediction to unify the gene prediction quality using Prodigal/2.6.3^33^ with the -a parameter to predict all potential genes. From these predicted proteins, we identified and aligned seven suggested marker genes using the tool ncldv_markersearch^31^ with the -c parameter to produce a multiple sequence alignment file by Clustal Omega. To limit the influence of non-informative sites, alignment columns containing more than 90% gaps were trimmed using trimAI/1.4.1^34^ with the -gt 0.1 parameter.

To minimize the influence of long branch attraction effects, we employed the posterior mean site frequency model, which necessitates a guide tree as input. We constructed this guide tree using IQ-TREE/2.2.0^35^, selecting the best-fit model by ModelFinder (-m MFP). The final phylogenetic tree was reconstructed under the parameters (-ft <guide tree> -m Q.pfam+F+I+I+R8+C60 -B 1000 -alrt 1000). For this, Q.pfam+F+I+I+R8 is the optimal model selected in the guide tree and the C60 matrix is used to implement the PMSF model^36^. -B and -alrt stands for the ultrafast bootstrap (UFboot) value and SH-aLRT test, respectively. The criterion that we used for the branch support was UFboot > 95% or SH-aLRT > 80% according to IQ-tree documentation. Both iTOL v6^37^ and anvi’o v7.1^38^ were used to visualize the viral tree.

### Reconstruction of gene trees

The predicted genes within the 195 viral genomes were grouped into OGs using OrthoFinder/2.5.4^39^ with -og parameters, yielding 8,876 OGs excluding singletons. Each OG was aligned using MAFFT/7.505, employing the E-INS-I model^40^ (--maxiterate 1000 -- genafpair). Alignment columns consisting of more than 90% gaps were trimmed using TrimAI/1.4.1 to reduce the influence of non-informative sites. These alignments were then subjected to maximum-likelihood gene tree reconstruction, leveraging the model that minimizes the Bayesian Information Criterion (BIC) via IQ-TREE/2.2.0 (-m MFP). During the tree reconstruction process, we documented 1,000 bootstrap trees (-wbtl) to be used for subsequent tree reconciliation analysis.

Of these 8,876 OGs, 5,285 OGs containing four or more genes were compared with the viral tree for evolutionary reconciliation by ALE. Among them, the tree reconciliation was successful for 4,786 OGs (see details below).

We identified viral-specific OGs among these 4,786 OGs through a sensitive homology search using Diamond/2.0.15^41^ (--very-sensitive). An OG was classified as viral-specific if none of its members matched any entry in the database, which comprised both KEGG (Kyoto Encyclopedia of Genes and Genomes) cellular organisms^42^ and metagenome-assembled genomes from marine planktonic eukaryotes, derived from the *Tara* Oceans project^43^, with an E-value threshold of 10^−3^. Consequently, we identified 3,535 of the 4,786 OGs as being viral-specific.

### Gene tree and viral tree reconciliation

The previously acquired viral tree and set of gene trees (the bootstrap trees for each OG rather than the consensus tree) were then subjected to the tree reconciliation tool ALE/0.4.0^27^ with the default parameters.

ALE is a probabilistic tool used to explore both gene-level and species-level events within a phylogenetic context. This method reconciles a collection of gene trees, such as a set of bootstrap trees from a gene family, with a predetermined species tree. Ancestral events, like originations, duplications, transfers, and losses, can thereby be inferred. Furthermore, it enables gene counting at every node of the species tree. ALE accounts for uncertainties of individual gene trees, which are often poorly resolved because of the low information carried in these genes. This approach calculates conditional clade probabilities, representing the weight of a certain gene tree topology from bootstrap samples (ALEobserve), and samples 100 reconciliations in the whole tree reconciliation space to avoid solely explaining the evolutionary scenario by the maximum likelihood reconciliation (ALEml_undated)^44^. We applied a threshold of 0.3 to the raw reconciliation frequencies produced by ALE, as suggested by previous studies^45,46^. Consequently, events such as loss, HGT, origination, or duplication, and the presence or absence of a gene, are recognized when they exhibit a frequency of at least 0.3 over internal nodes or tips. This threshold is thought to enhance the sensitivity of the analysis, permitting even low-frequency events to be detected.

When an OG is reconciled with a viral tree, a maximum likelihood value of the loss rate is estimated. Reconciling order-specific OGs with the complete viral tree can possibly increase the loss rate, consequently influencing the inference of other evolutionary events. To minimize this effect, we assumed that order-specific OGs were gained at the root of the order or during the evolution of the order and were therefore analyzed separately from other OGs. For order-specific OGs, we used the subtree corresponding to the viral order instead of using the whole viral tree for tree reconciliation. For other OGs, we used the whole viral tree as the reference for tree reconciliation.

For some OGs (n = 499), ALE reconciliations were unsuccessful. These fell into two cases: 1) they lacked sufficient informative genes within the gene family to conduct a bootstrap test, or 2) after trimming, they contained too many identical sequences, resulting in a non-bifurcated gene tree that hindered subsequent reconciliation.

### Analysis of evolutionary events rate

RED^29^, which forms the basis for analyzing the dynamics of the evolutionary event rate, was computed using the “get_reds” function of the “castor” R package^47^. The normalized evolutionary rate was then determined by taking the natural logarithm (plus one) of the number of events at each node divided by the corresponding branch length leading to that node. We performed the Mann-Whitney U test using the R package “ggsignif”^48^ to examine the rate differences of the various evolutionary events. Average nucleotide identity (ANI) was calculated by fastani/1.33^49^.

## Acknowledgements

We would like to thank Russell Y. Neches for his critical comments on this work. This work was supported by JSPS/KAKENHI (18H02279 and 22H00384, to H.O.), JST SPRING, Grant Number JPMJSP2110, and the Collaborative International Joint Research Program of the Institute for Chemical Research, Kyoto University (No. 2022-26 to M.G.). Computational time was provided by the SuperComputer System, Institute for Chemical Research, Kyoto University. We thank J. Iacona, Ph.D., from Edanz (https://jp.edanz.com/ac) for editing a draft of this manuscript.

## Author contributions

J.W. and H.O. designed the study. J.W. performed all bioinformatics analysis. L.M. contributed to the preparation of datasets and helped with the bioinformatics analyses. H.O. supervised the work and M.G., H.H., Y.O., and H.E. co-supervised. J.W. wrote the initial draft of the manuscript. All authors contributed to data interpretation and approved the final version of the manuscript.

