## Supplementary Material for "Gene transfer among viruses substantially contributes to gene gain of giant viruses"

**Table S1. Examples of evolutionary events summarized from root to individual virus.**

| Viruses | Order | Genes for Reconciliation | Total Gains | Total Losses | Log(Gains / Losses) |
|---|---|---|---|---|---|
| Paramecium bursaria Chlorella virus NY2A | Algavirales | 346 | 545 | 204 | 0.9826658 |
| Ostreococcus tauri virus OtV5* | Algavirales | 202 | 424 | 243 | 0.5566720 |
| Kaumoebavirus strain KLCC10 | Asfuvirales | 224 | 273 | 28 | 2.2772673 |
| Abalone asfarvirus* | Asfuvirales | 55 | 128 | 69 | 0.6179238 |
| Melanoplus sanguinipes entomopoxvirus | Chitovirales | 187 | 239 | 38 | 1.8388774 |
| Cotia virus SPAn232* | Chitovirales | 157 | 365 | 199 | 0.6065925 |
| Tupanvirus soda lake | Imitervirales | 770 | 1120 | 263 | 1.4489299 |
| Namao virus* | Imitervirales | 110 | 354 | 270 | 0.2708750 |
| Emiliania huxleyi virus 202 | Pandoravirales | 395 | 563 | 143 | 1.3704350 |
| Mollivirus sibericum* | Pandoravirales | 165 | 327 | 200 | 0.4916428 |
| Orpheovirus IHUMI-LCC2 | Pimascovirales | 557 | 693 | 140 | 1.5993876 |
| Dikerogammarus haemobaphes virus 1 | Pimascovirales | 11 | 178 | 201 | -0.1215214 |
| Lymphocystis disease virus 1 | Pimascovirales | 56 | 239 | 218 | 0.0919685 |

This table presents three selected examples (the highest, lowest, and closest to zero) from each viral order using the log-transformed ratio of gene gain events to gene loss events. A positive log-transformed value indicates a tendency of gene gains over losses during the evolution of each virus. Conversely, a negative value signifies a higher frequency of gene losses than gains. A value of zero signifies an equivalent rate of gene gain and loss during evolution. A single asterisk (*) denotes the lowest value and the value closest to zero pertaining to the same virus.

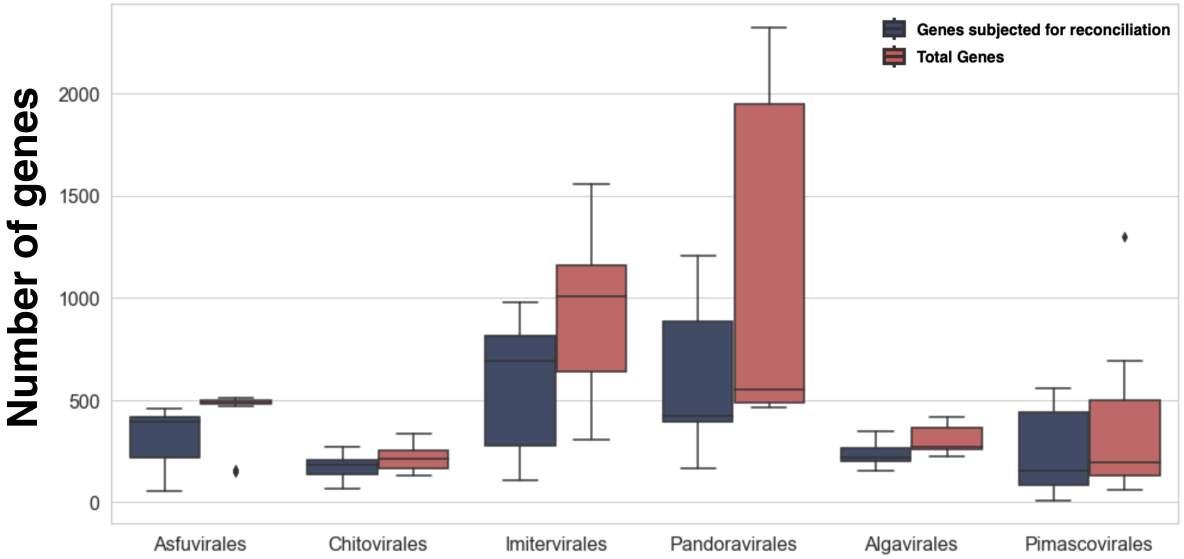


**Figure S1. Comparison of the total gene count and number of genes subjected to tree reconciliation analysis.**

The boxplot presents comparisons between the number of genes encoded in the individual viral genomes and number of genes that were used for tree reconciliation analysis.

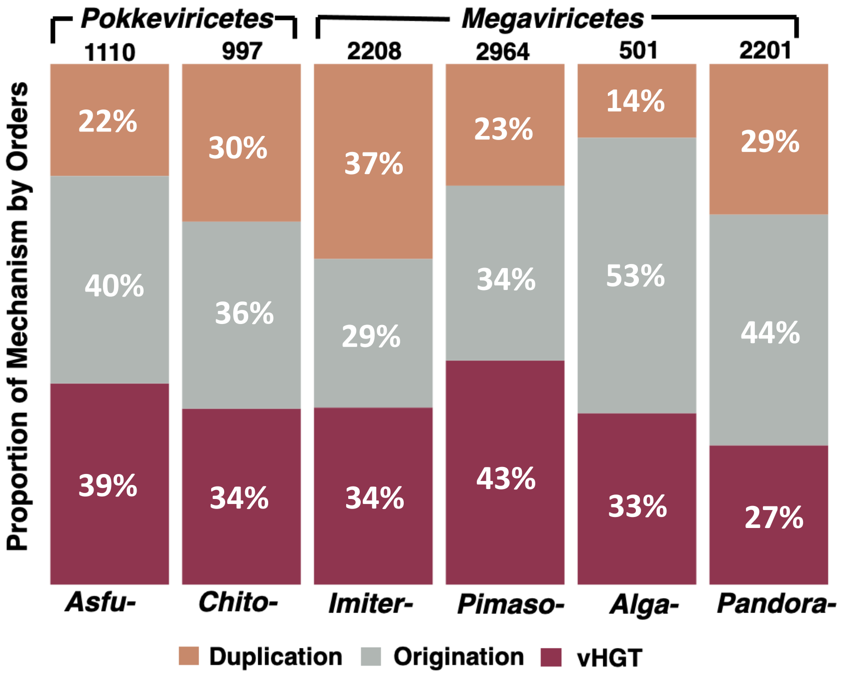


**Figure S2. Comparative contributions of various gene gain mechanisms across viral orders within virus-specific OGs.**

The elements in this plot mirror those of Figure 3.

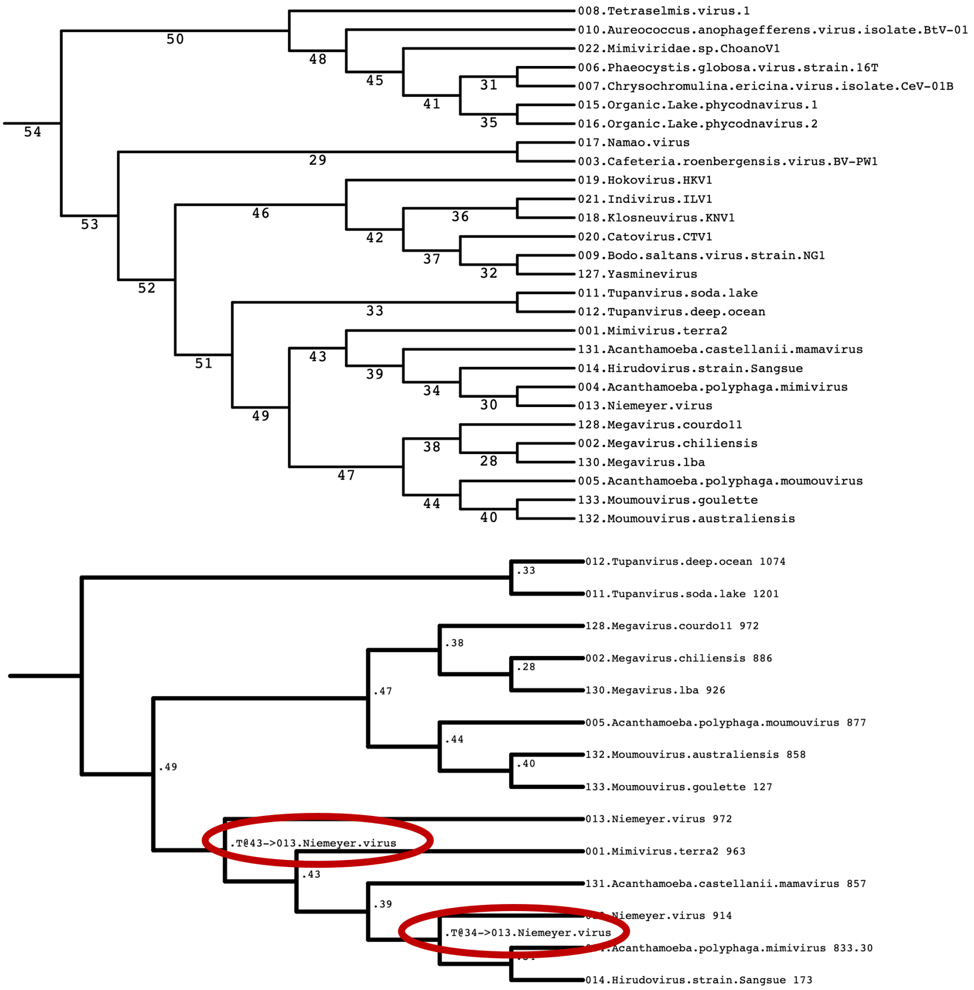


**Figure S3. Example of visualization the tree reconciliation results.**

We selected the tree reconciliation results of OG0001042 as a representative example of visualizing detected vHGT. Two vHGT events were inferred to explain the existence of two copies of a gene in Niemeyer virus. The upper tree is the subtree that contains viruses from the *Imitervirales* order, and the numbers represent the ID of internal nodes. The lower tree is one of the sampled reconciliation gene tree topologies with annotated events that are consistent with the final results (explained two vHGT events). In this tree, the texts beside the internal nodes represent evolutionary events. For example, ".33" indicates the evolution of this gene in two tupanviruses following their speciation event (as indicated with "33" in the upper viral tree). The text ".T@43 -> 013.Niemeyer.virus" indicates that there was a vHGT from internal node "43" to Niemeyer virus (leaf "013" in the viral tree). In this case, it is plausible that an unsampled virus branching out from near node "43" was the doner for this vHGT.

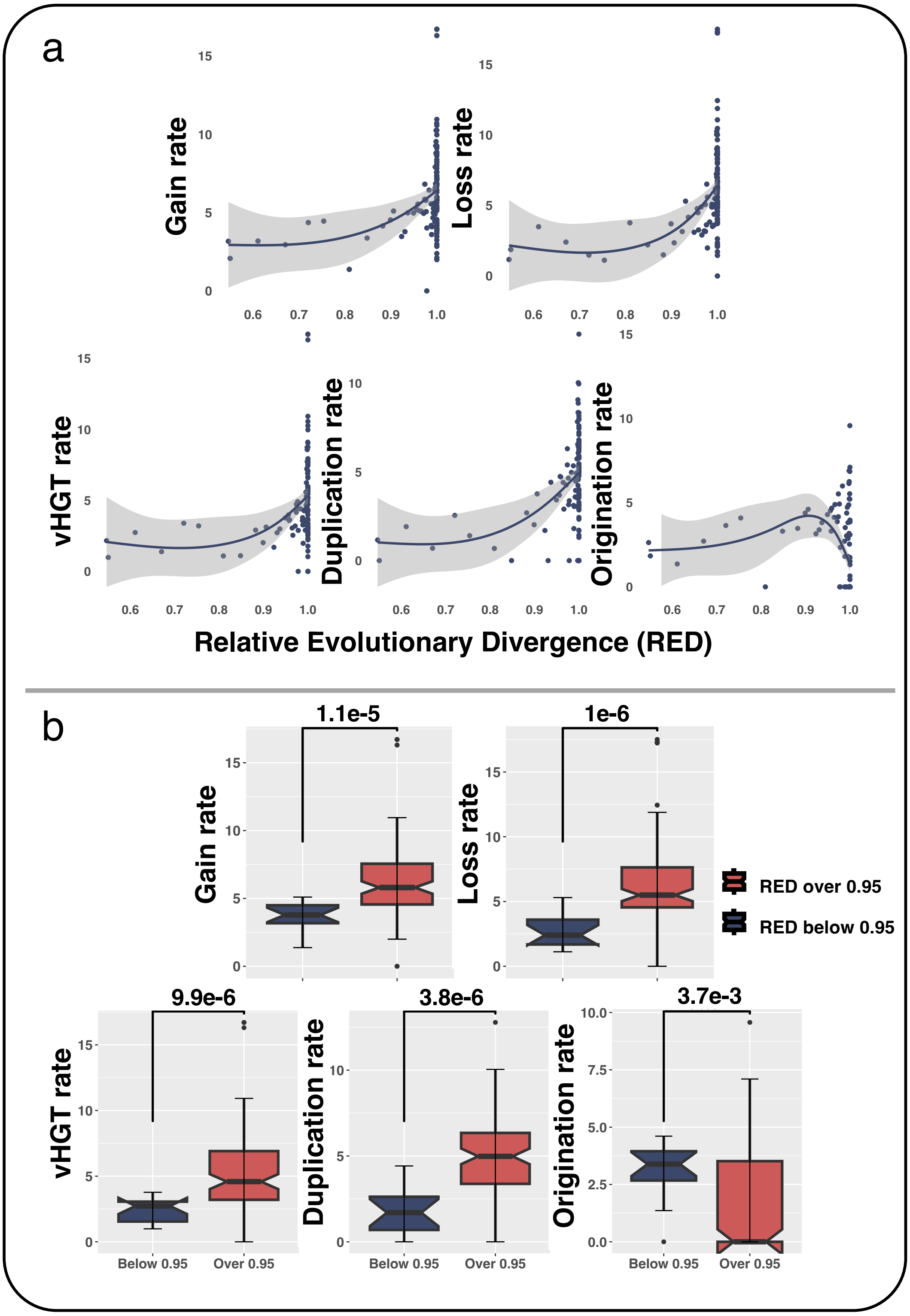


**Figure S4. Rates of different evolutionary events along the divergence of *Pokkeviricetes*.** (a) The evolutionary rates of different evolutionary events are plotted against the divergence measured by RED. (b) Boxplot provides a comparison of evolutionary rates between recent (RED ≥ 0.95) and earlier periods (RED < 0.95). *P*-values from the Mann-Whitney U test are shown above the graph.

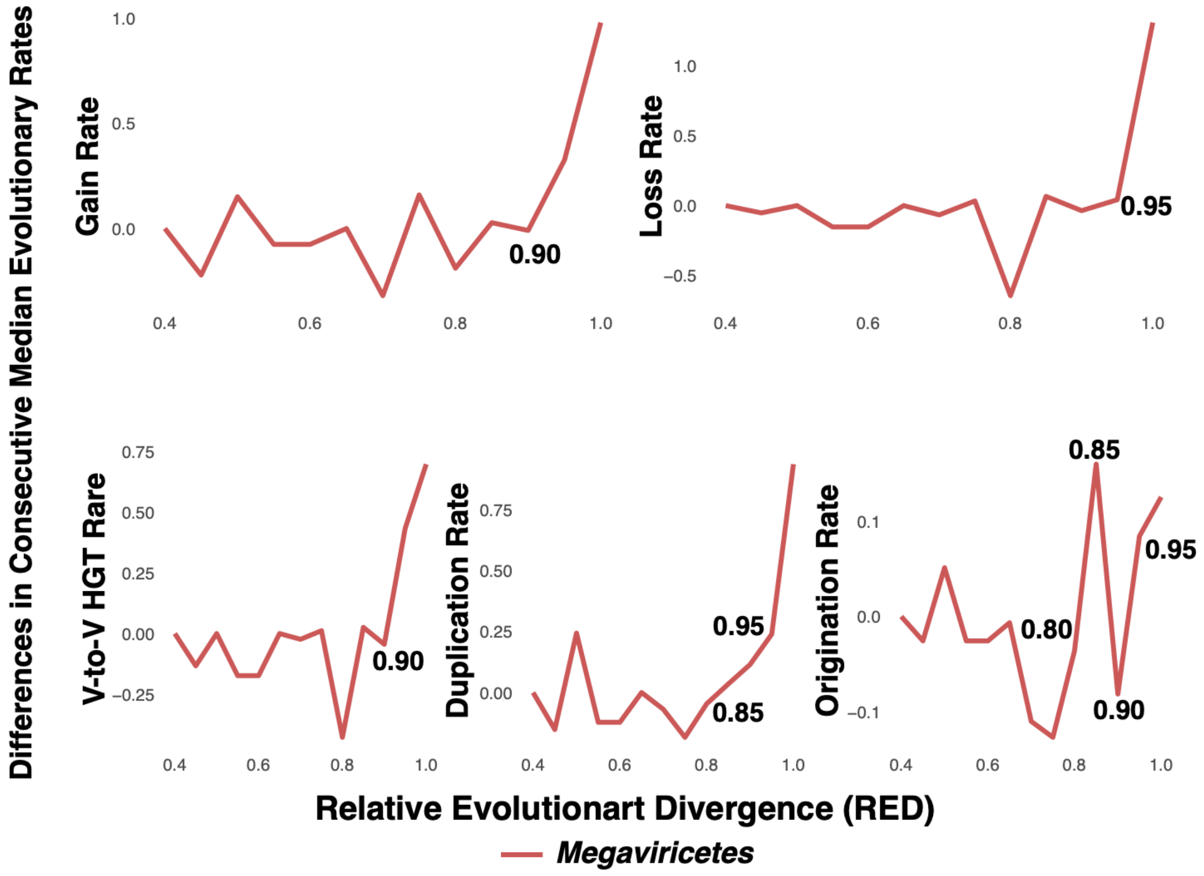


**Figure S5. Changes of evolutionary rates within *Megaviricetes*.**

Changes in the median rates of evolutionary events within the *Megaviricetes* lineage over time are shown. First, the median rate at a Relative Evolutionary Distance (RED) of x is computed using data within the range (0, x). Then, the median rate at a previous time step (RED = x - 0.05) is subtracted from the median rate at RED = x. This difference is plotted along the RED value by setting the first value to RED = 0.4 to 0. Notable positive changes in the evolutionary event rate are observed during recent evolution. We set RED = 0.95 as the boundary of the rate change for comparative analyses shown in Figure 4.

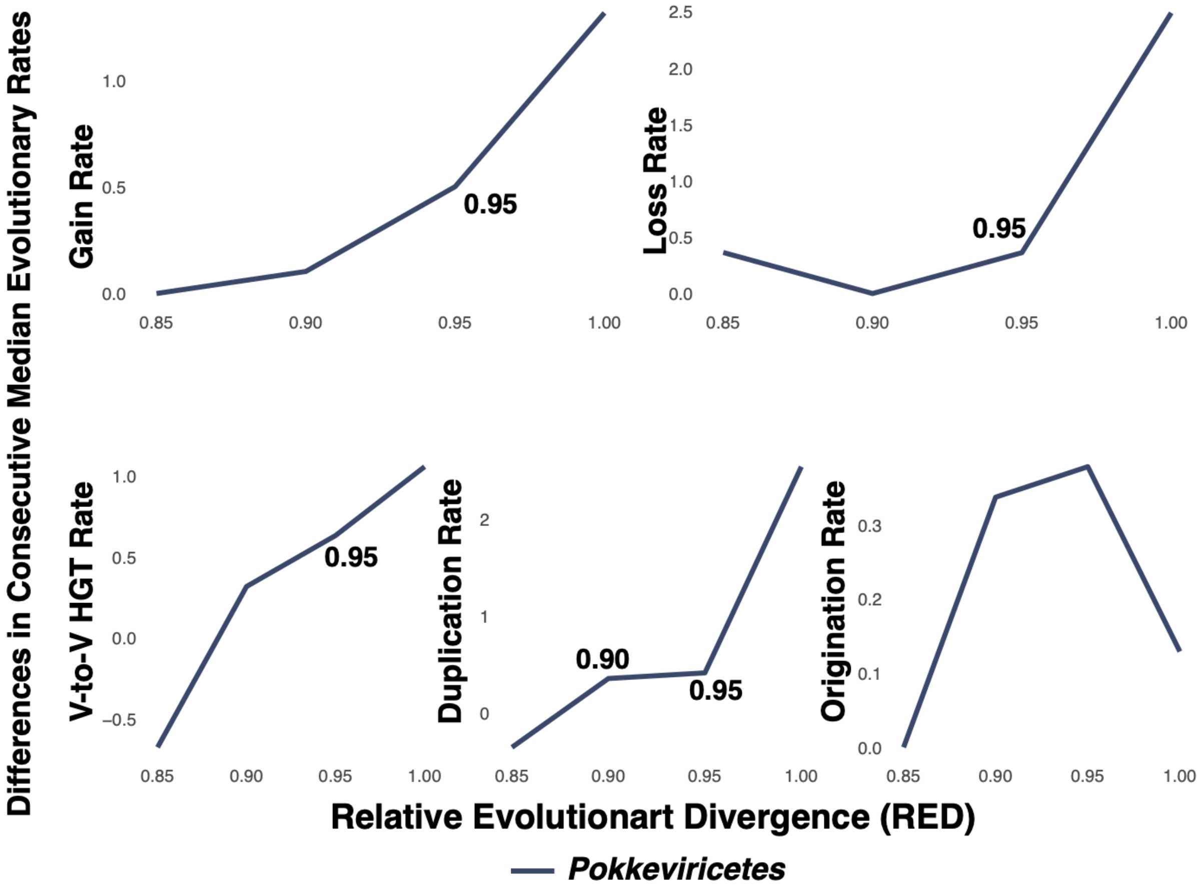


**Figure S6. Changes of evolutionary rates within *Pokkeviricetes*.**
Changes in the median rates of evolutionary events within the *Pokkeviricetes* lineage over time are shown. First, the median rate at a Relative Evolutionary Distance (RED) of x is computed using data within the range (0, x). Then, the median rate at a previous time step (RED = x - 0.05) is subtracted from the median rate at RED = x. This difference is plotted along the RED value, by setting the first value at RED = 0.4 to 0. Notable positive changes in the evolutionary event rate are observed during recent evolution. We set RED = 0.95 as the boundary of the rate change for comparative analyses shown in Figure S3.
